# CPPLocPred: Subcellular Localization of Cell-Penetrating Peptides

**DOI:** 10.64898/2026.07.28.741285

**Authors:** Nisha Bajiya, Naman Kumar Mehta, Gajendra P. S. Raghava

## Abstract

Cell-penetrating peptides (CPPs) are widely used to deliver therapeutic cargoes into cells. Although numerous computational methods have been developed for identifying CPPs and several predictors are available for protein subcellular localization, no method has been developed to predict the subcellular localization of CPPs. Here, we present CPPLocPred, a hierarchical machine-learning (ML) framework that predicts CPPs and their subcellular localization. In the first stage, we developed ML models to identify CPPs, achieving an AUC of 0.953 with an MCC of 0.7842 on an independent set, exhibiting performance equivalent to or better than existing state-of-the-art methods. In the second stage, we developed a method for predicting the subcellular localization of CPPs. Subcellular localization methods were trained (80% data using five-fold cross-validation) and validated (20% data) on experimentally validated CPPs for 663 Cytoplasm, 287 Nucleus, 57 Mitochondria, 186 Endo_lysosome, and 328 Others. Our primary analysis revealed that Mitochondrial and Nuclear associated CPPs are abundant in positively charged arginine- and lysine-rich patterns, whereas Endo_lysosomal CPPs preferentially comprise glycine-, proline-, and cysteine-rich motifs. We used a wide range of traditional peptide features, along with the embedding of protein language models, to develop ML models. Among all evaluated models, the CatBoost-based subcellular localization models with Distance Distribution of Residues (DDR) achieved AUCs of 0.814, 0.775, 0.970, 0.782, and 0.798 for Cytoplasm, Nucleus, Mitochondria, Endo_lysosome, and Others, respectively, on validation dataset. We developed CPPLocPred, which offers a practical platform for functional annotation and rational design of localization-specific CPPs for therapeutic applications (https://webs.iiitd.edu.in/raghava/cpplocpred/).

**Highlights:**

- Prediction and subcellular localization of cell-penetrating peptides.
- Localization of CPPs depends on their amino acid and dipeptide composition.
- Best feature for subcellular localization was Distance Distribution of Residues.
- CatBoost model achieved the highest performance for subcellular localization.
- A web server and standalone software to facilitate its use by the scientific community.

## Introduction

Over the past two decades, drug discovery has witnessed a significant shift from small-molecule drugs toward peptide- and protein-based therapeutics, as reflected by the growing number of FDA-approved peptide and protein drugs [1][2][3][4][5][6][7]. Therefore, researchers are developing computational methods and databases for a wide range of bioactive peptides, including anticancer, antibacterial, antiparasitic, anti-angiogenic, toxic, hemolytic, and cell-penetrating peptides [8][9][10][11][12][13]. Cell-penetrating peptides (CPPs) are a diverse class of short peptides (typically 5-30 amino acids) that can traverse cell membranes and deliver various molecules, such as peptides, proteins, nucleic acids, small-molecule drugs, and nanoparticles [14]. Since the discovery of the transactivator of transcription (TAT) peptide from HIV-1 and penetratin from the Antennapedia homeodomain, CPPs have emerged as promising delivery vehicles for overcoming one of the major challenges in drug development: efficient transport across cellular membranes [15][16]. Their increased intracellular uptake and low toxicity have led to CPPs being utilized for cancer therapy, gene delivery, vaccination and the development of antimicrobial drugs and targeted drug delivery systems [17].

Although substantial progress has been made in CPP research, successful intracellular delivery involves more than simply crossing the plasma membrane. Following cellular uptake, CPPs and their cargoes must reach specific intracellular compartments to exert their intended biological functions. For example, peptides designed for gene regulation or genome editing require efficient nuclear localization, whereas peptides targeting mitochondrial dysfunction must accumulate within mitochondria. Similarly, retention within endosomal or lysosomal compartments can significantly influence delivery efficiency and therapeutic outcome [18][19][20]. Consequently, intracellular localization is a critical determinant of CPP functionality and therapeutic success.

Experimental determination of CPP localization remains labor-intensive, time-consuming, and expensive. As a result, computational approaches for accelerating CPP discovery and characterization have become increasingly investigated. Over the past decade, several machine-learning (ML) methods have been developed for CPP classification and prediction of uptake efficiency (Table 1). In addition to conventional sequence-derived features, physicochemical descriptors, and ensemble-learning approaches were also employed. More recently, protein language models (PLMs) such as ProtBERT and ESM2 have achieved even higher prediction accuracy by directly capturing contextual, structural, and evolutionary information from protein sequences [21][22].

**Table 1:**
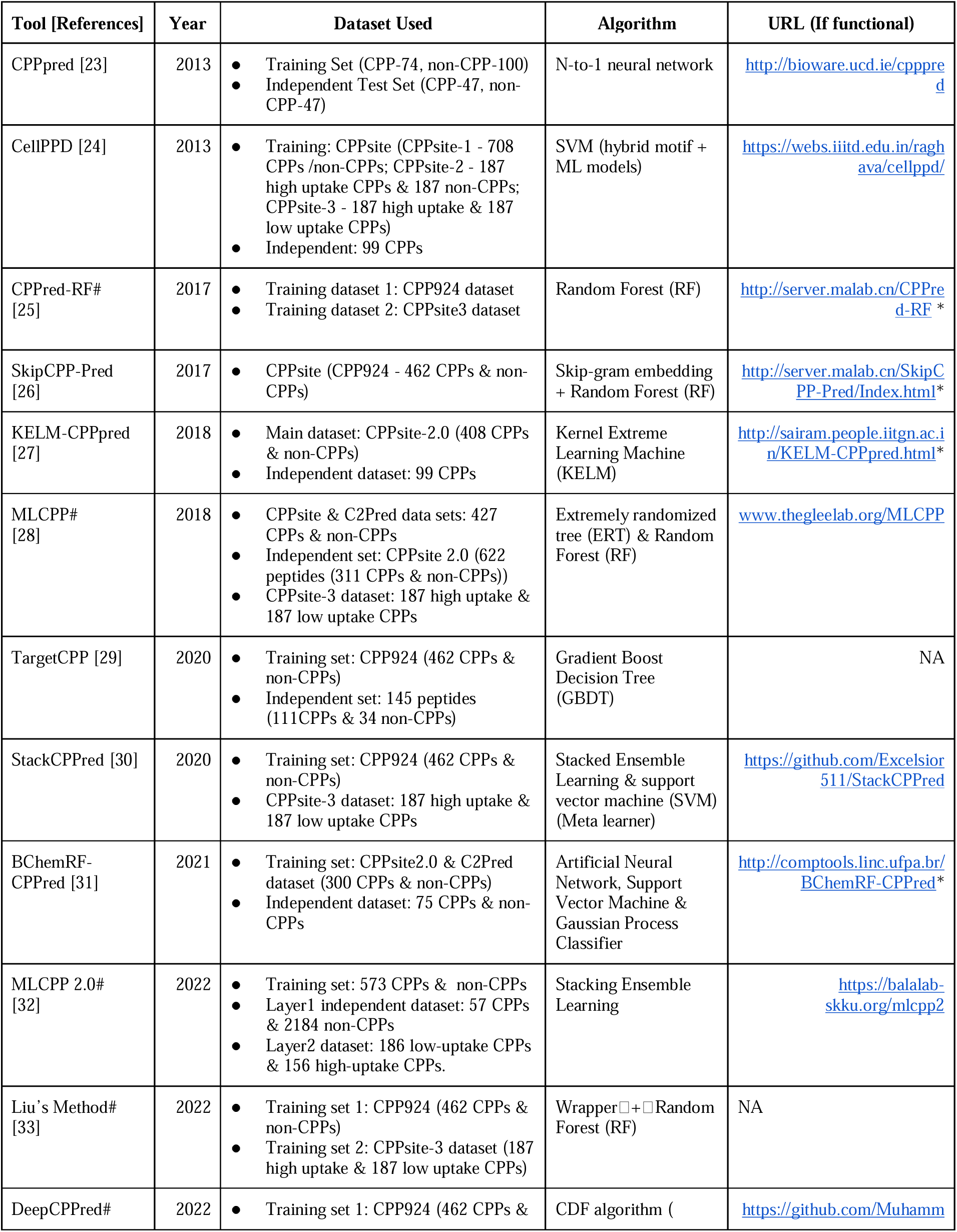
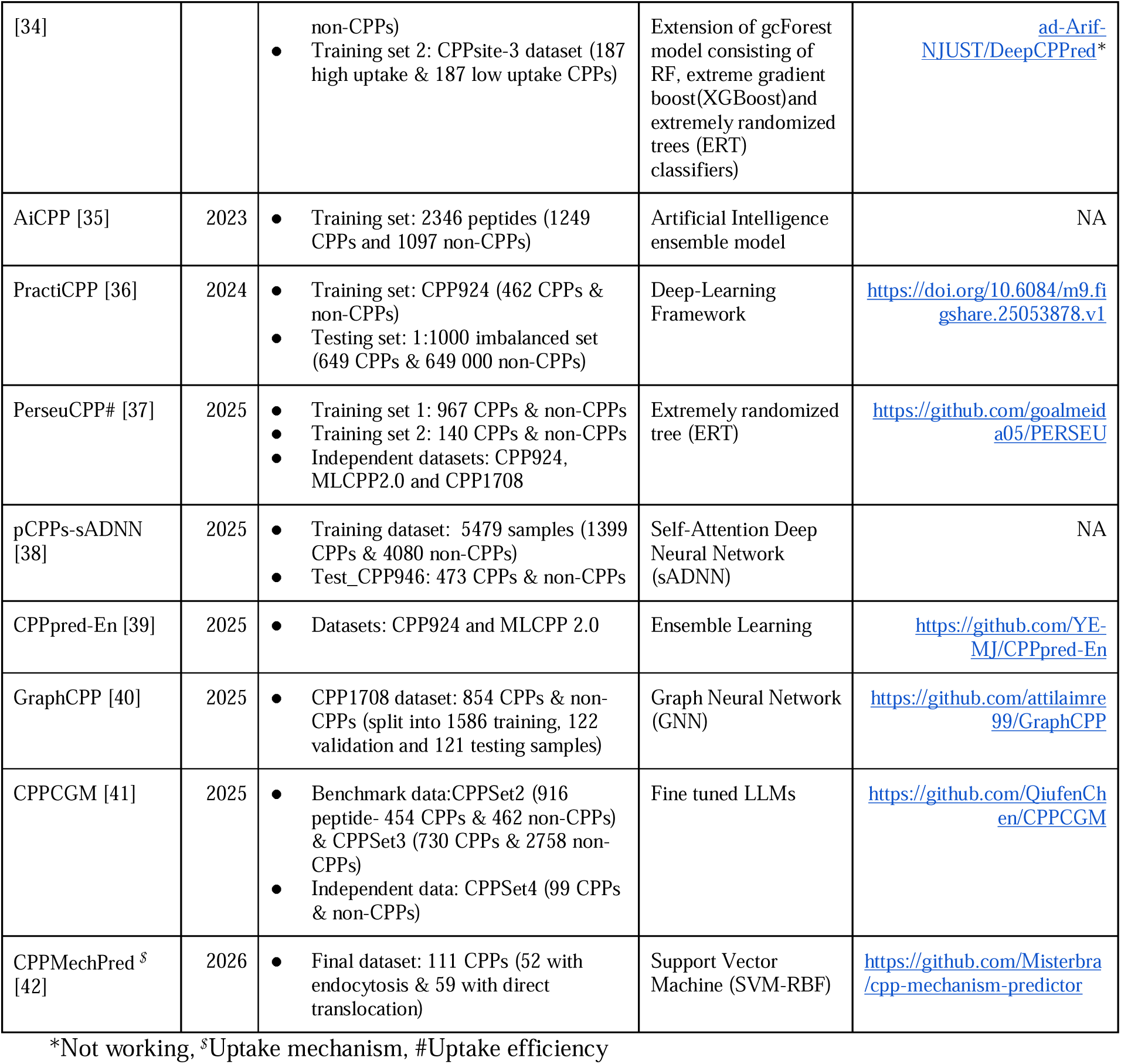
List of tools developed for predicting cell-penetrating peptides.

Despite these advances, existing methods only predict whether it will penetrate inside the cell or not. These methods are suitable for identifying CPPs that can deliver cargo inside a cell, but are not suitable for delivering cargo to a specified location within a cell. Notably, subcellular localization prediction has been studied extensively for proteins [43][44], with several well-established computational tools available for this purpose [45][46][47][48][49][50], yet no comparable prediction tool currently exists for peptides. In this regard, it is important to emphasize that peptides having a similar ability for cell penetration may have very different behaviors in terms of intracellular transport and biological functions, highlighting a significant knowledge gap. Furthermore, distinct subcellular compartments are associated with specific sequence signatures, physicochemical properties, and targeting motifs. For example, nuclear localization signals (NLS) are defined by motifs rich in lysine and arginine that are specifically recognized by nuclear importins, while mitochondrial targeting peptides (MTPs) are generally enriched in positively charged arginine residues and depleted in acidic amino acids [51][52]. Thus, understanding the sequence determinants of these targeted organelles could be used to design rational delivery strategies to a specific subcellular compartment.

The recent release of CPPsite3 [53] has expanded the collection of experimentally verified CPPs, accompanied by useful annotations on subcellular localization and peptide characteristics that are valuable in pursuing and building predictive models of CPPs’ subcellular targeting. In the present study, we present a hierarchical computational framework, CPPLocPred, designed for simultaneously predicting CPPs and their subcellular localization (Figure 1). First, experimentally validated CPPs were analyzed to identify localization-associated residue composition patterns, dipeptide signatures, physicochemical properties, positional residue preferences, and conserved motifs. Subsequently, multiple traditional descriptors, PLM embeddings, ML and DL algorithms, feature-fusion strategies, and motif-based approaches were systematically evaluated. Based on extensive benchmarking, a prediction framework was established that first identifies CPPs from non-CPPs and, second, predicts their subcellular location from one of five major categories: Cytoplasm, Mitochondria, Nucleus, Endo_lysosome, and Others. Finally, the best-performing models were implemented in a freely accessible web server, CPPLocPred, to facilitate large-scale prediction and rational design of localization-specific CPPs.

**Figure 1:**
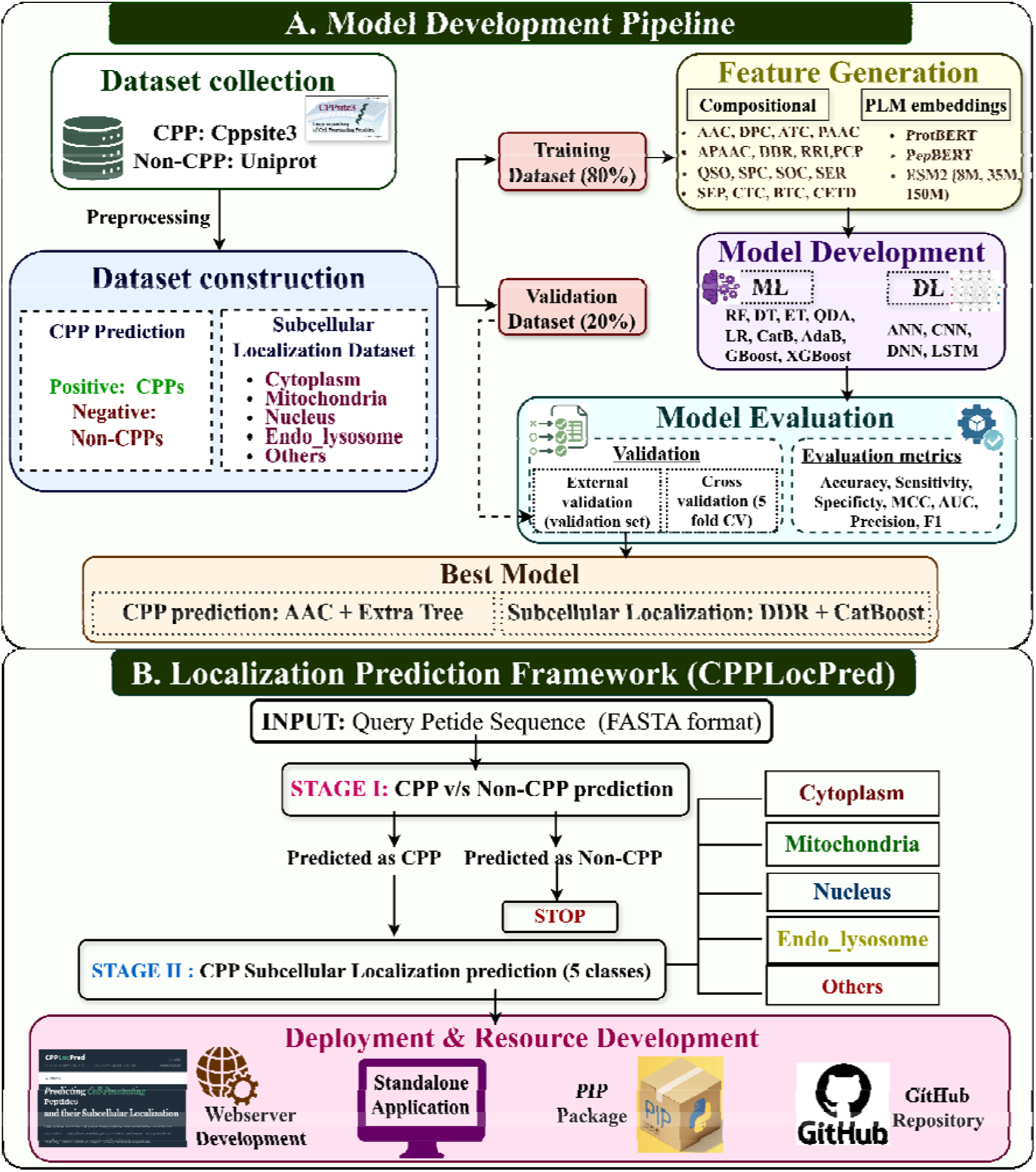
The complete workflow of CPPLocPred.

## Materials and Methods

### 1. Dataset Collection and Preparation

Experimentally validated CPPs were obtained from the latest version of CPPsite3 [53], which has over 6,000 CPPs annotated with various biological features. To construct the localization dataset, CPPs were categorized into five broad subcellular localization classes with several subcategories: Cytoplasm, Nucleus, Mitochondria, Endo_lysosome, and Others. The Endo_lysosome class included peptides localized to endosomes, lysosomes, and vesicles, whereas the Others class comprised peptides found in membranes, vacuoles, Golgi bodies, periplasm, plastids, and other intracellular structures. To ensure dataset quality, peptides containing modified, cyclic, or attached side-linker residues, D-amino acids, unnatural amino acids, or additional chemical groups were excluded. Only linear peptides composed of natural L-amino acids were retained. Duplicate sequences were subsequently removed, resulting in 703 Cytoplasmic, 287 Nuclear, 186 Endo_lysosomal, 57 Mitochondrial, and 328 Others CPPs with sequence lengths ranging from 5 to 50 residues. Detailed sequence counts at each filtering step are provided in Supplementary Table S1.1 and S1.2.

### 2. Construction of Prediction Datasets

#### 2.1. CPP Prediction

For CPP classification, all localization specific CPP datasets were merged, and duplicate sequences were removed, yielding a final positive dataset of 1,366 unique CPPs. To generate a high-quality negative dataset, peptide fragments were generated from downloaded UniProt protein sequences with the keyword “Proteins”, which are reviewed and range in length from 1 to 200 residues. Peptides corresponding to known CPPs were removed, and an in-house Python script was used to eliminate sequences sharing ≥90% sequence identity with positive CPPs of the same length. The remaining peptides were randomly sampled to match the length distribution of the positive dataset, resulting in a balanced dataset containing 1,366 CPPs and 1,366 non-CPPs (Table 2). To assess the generalizability of our CPP classifier, we created a separate independent dataset by taking positive sequences from CPPsite3 whose location was unknown and then applying the same filters as for our main dataset creation for modification removal, yielding 810 experimentally validated CPPs ranging from 5 to 50 residues that were not used for training and validation. Additionally, we deleted duplicate sequences from our independent set that were included in the existing datasets used to create the CPP prediction models, resulting in a final count of 758 positive sequences. Similarly, 758 negative sequences were randomly generated using an in-house Python script, and CD-HIT [54] was applied at various thresholds to reduce redundancy.

**Table 2:** Number of peptides in datasets used for prediction and subcellular localization.

| Category | Positive set | Negative set | Total |
| --- | --- | --- | --- |
| **CPPs Prediction** |  |  |  |
| **CPPs and Non-CPPs** | 1366 | 1366 | 2732 |
| **Subcellular Localization Prediction** |  |  |  |
| **Cytoplasm** | 663 | 663 | 1326 |
| **Nucleus** | 287 | 287 | 574 |
| **Mitochondria** | 57 | 57 | 114 |
| **Endo_lyosome** | 186 | 186 | 372 |
| **Others** | 328 | 328 | 656 |

#### 2.2. CPP Subcellular Localization

For subcellular localization prediction, a one-versus-rest (OvR) strategy was adopted. For each localization class, peptides belonging to the target location were considered positive samples, whereas peptides from all remaining locations served as negative samples. To prevent information leakage, sequences present in the positive dataset were omitted from the corresponding negative dataset. To eliminate ambiguity during training, sequences appearing in multiple compartments were excluded. Because the localization data exhibited considerable class imbalance, particularly for Mitochondrial and Endo_lysosomal peptides, balanced datasets were generated by randomly selecting an equal number of negative samples from the remaining classes while maintaining peptide length distributions. For the Cytoplasmic class, where the positive set (703 peptides) exceeded the available negative set (663 peptides), 663 positive sequences were randomly selected to maintain class balance. The final 5 localization datasets (Table 2) were employed to develop separate binary classifiers for each class. Eventually, the overall prediction framework adopted a hierarchical architecture comprising two stages. At first, peptides were classified as CPPs or non-CPPs. Only peptides predicted as CPPs were forwarded to localization-specific classifiers to predict their subcellular locations.

### 3. Cross-validation approach

To ensure unbiased model evaluation, datasets were split into training (80%) and validation (20%) sets using stratified sampling, preserving the original class distributions. As shown in Figure 2, the training set consisted of 1,061 Cytoplasmic, 459 Nuclear, 91 Mitochondrial, 298 Endo_lysosomal, and 525 Others CPPs. The validation set contained 608 sequences, including 265 “Cytoplasmic”, 115 “Nuclear”, 74 “Endo_lysosomal”, 23 “Mitochondrial”, and 131 “Others” CPPs. The models were built, and hyperparameters were tuned exclusively on the training data through five-fold stratified cross-validation (CV). In each fold, four subsets were used for training, and one for testing. This process was repeated 5 times so that each subset acted as the test data at once, following the conventional strategy. The validation data were left completely unrevealed during the model training, feature optimization and tuning and was utilized solely for final model evaluation.

**Figure 2:**
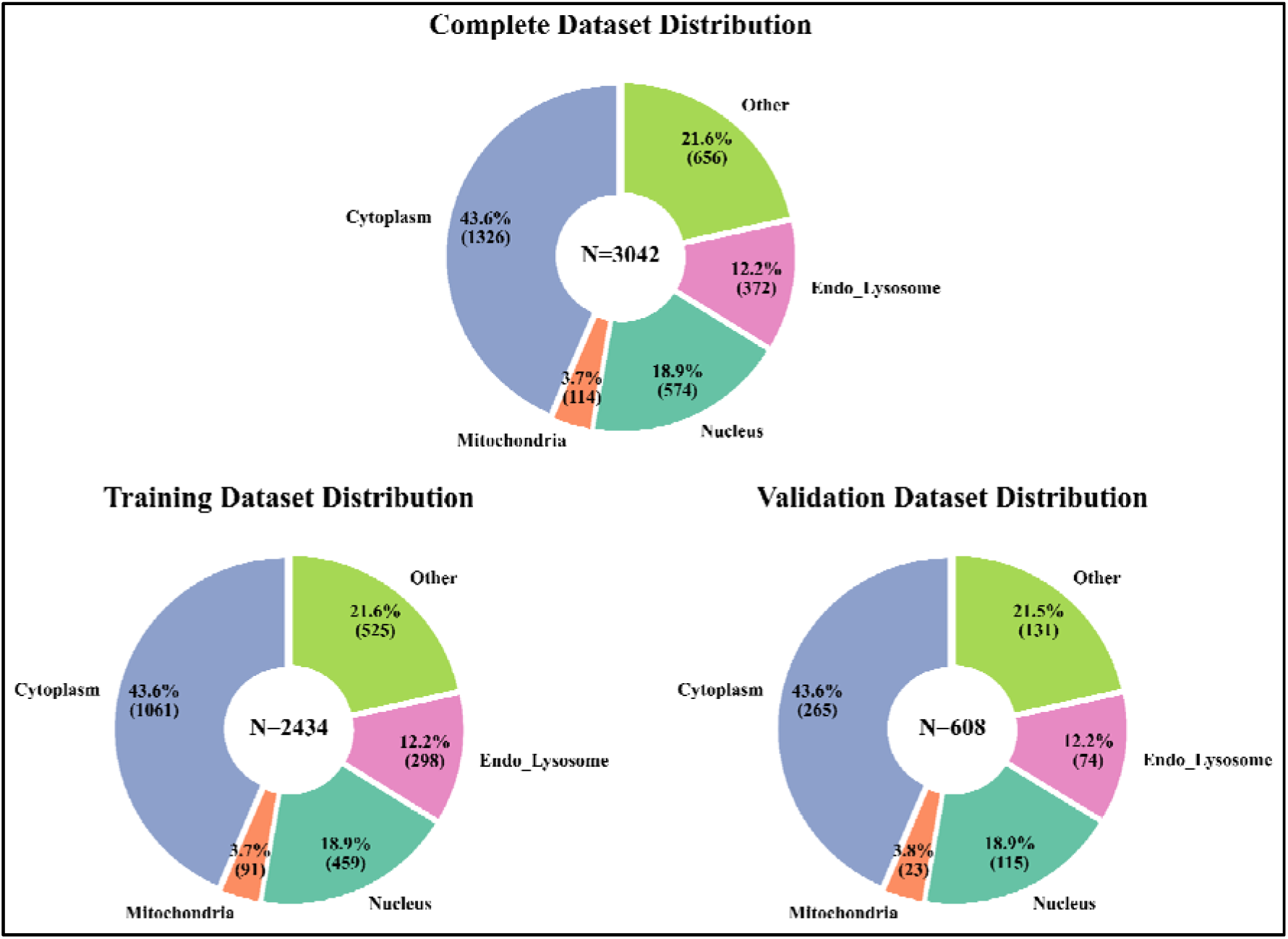
Dataset distribution of CPP localization classes.

### 4. Preliminary Sequence Analysis

#### 4.1. Amino Acid Composition

The amino acid composition (AAC) of each peptide was calculated as the percentage occurrence of individual amino acids within the sequence. For each amino acid residue (i), the composition was calculated in Equation 1.

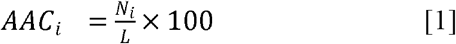

where *N_i_* represents the number of occurrences of residue *i,* and *L* denotes the peptide length. Mean amino acid compositions were subsequently calculated for each localization class and the non-CPP dataset.

#### 4.2. Dipeptide Composition

Dipeptide composition (DPC) was calculated for all 400 possible dipeptide combinations to capture local sequence-order information. For a given dipeptide (ij), the composition was computed as Equation 2:

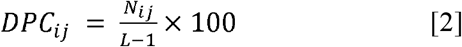

Where *N_ij_* is the frequency of dipeptide *ij* in the peptide sequence, and *L* represents the peptide length. Mean dipeptide compositions were subsequently determined for each localization class and the non-CPP dataset.

#### 4.3. Physicochemical Property

A range of physicochemical properties was calculated using the ProteinAnalysis module implemented in <u>Biopython</u> [55], including peptide length, molecular weight, aromaticity, instability index, isoelectric point (pI), grand average hydropathy (GRAVY), and net charge. In addition, residue-group compositions were calculated manually as percentages of residues belonging to six physicochemical categories: Basic (K, R, H), Acidic (D, E), Polar (S, T, N, Q, C, Y), Nonpolar (A, V, L, I, M, F, W, P, G), Aliphatic (A, V, I, L), and Neutral residues (all residues excluding acidic and basic amino acids) [56]. The composition of each residue group was calculated as in Equation 3.

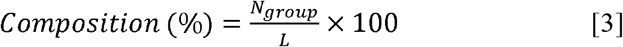

where *N_group_* represents the number of residues belonging to a given physicochemical category, and *L* denotes the peptide length. Hydrophilicity was calculated as the negative of the GRAVY score.

#### 4.4. Position-Specific Residue Analysis

Two-Sample Logo (TSL) analysis was performed to determine specific amino acids preference with the position associated with CPP localization [57]. Peptide sequences were converted into a pattern of fixed length of 10 residues by selecting 5 residues from each terminal (since the minimum length in our dataset was 5). Residue enrichment and depletion at each position were evaluated using the binomial test implemented in the TSL program. Amino acids significantly enriched in the positive dataset were displayed above the baseline, whereas depleted residues were displayed below the baseline. The height of entire stacks demonstrates how conserved each position is in terms of sequence, while the height of each symbol reflects the specific amino acids’ relative frequency at the position.

### 5. Alignment-based approach using Motif Analysis

Motifs are short amino acid patterns known to play a significant role in the subcellular localization of proteins and peptides inside cells. Localization-specific sequence motifs were identified using MERCI (Motif-EmeRging and with Classes-Identification) [58], a conserved motif identification method that uses a Perl script with various parameter combinations. Both the positive and negative training datasets are provided as inputs, and the tool identifies motifs that occur exclusively or preferentially in one class relative to another, allowing it to differentiate between localization classes. To capture both precise sequence motifs and generalized physicochemical patterns, four amino acid classification schemes were used: NONE, BETTS-RUSSELL, KOOLMAN-ROHM, and RASMOL. Multiple combinations of motif length (-l), motif count (-k), and minimum positive sequence frequency (-fp) were tested to determine the best parameter settings. The final parameter combination was chosen based on prediction accuracy. Identified motifs were sorted based on the number of positive hits obtained in the validation dataset and were subsequently used to interpret localization-specific sequence signatures and ML predictions.

### 6. Computing Peptide Features

To generate informative numerical representations of peptide sequences, both traditional sequence-derived descriptors and PLM embeddings were explored.

#### 6.1. Compositional Features

Sequence-based descriptors were generated using the pfeature package [59]. The evaluated descriptors included Amino Acid Composition (AAC), Dipeptide Composition (DPC), Amphiphilic Pseudo Amino Acid Composition (APAAC), Pseudo Amino Acid Composition (PAAC), Residue Repeat Information (RRI), Distance Distribution of Residues (DDR), Composition Enhanced Transition Distribution (CETD), Shannon Entropy-based descriptors (SEP, SER, and SPC), Quasi Sequence Order (QSO), and Sequence Order Coupling (SOC). Each descriptor was independently evaluated for its ability to discriminate CPPs and predict subcellular localization.

#### 6.2. Protein Language Model Embeddings

To capture contextual and evolutionary information from peptide sequences, embeddings were generated using five pretrained PLMs: ProtBERT (ProtBERT-BFD), PepBERT, ESM2-8M (ESM2_t6_8M_UR50D), ESM2-35M (ESM2_t12_35M_UR50D), and ESM2-150M (ESM2_t30_150M_UR50D). Peptide sequences were provided as raw amino acid sequences and converted into fixed-length embedding vectors using pretrained model weights without additional fine-tuning. Rostlab’s ProtTrans framework [21], was used to construct ProtBERT embeddings, whereas PepBERT (PepBERT-large-UniParc) is a peptide-specific language model pretrained on large-scale peptide datasets for peptide-related prediction tasks [60]. ESM2 models were obtained from Meta AI and pretrained on the UniRef50 database, allowing for the extraction of contextual, evolutionary, and structural sequence information [22]. Table 3 shows the generated embeddings, which were subsequently used as input features for localization prediction models.

**Table 3:** List of pretrained PLM employed for peptide representation and embedding generation.

| Embedding Name | Model Family | Full Model Name | Parameters | Training Data | Dimension |
| --- | --- | --- | --- | --- | --- |
| ESM2-8M | ESM2 | ESM2_t6_8M_UR50D | 8 Million | UniRef50 | 320 |
| ESM2-35M | ESM2 | ESM2_t12_35M_UR50D | 35 Million | UniRef50 | 480 |
| ESM2-150M | ESM2 | ESM2_t30_150M_UR50D | 150 Million | UniRef50 | 640 |
| ProtBERT | ProtTrans | ProtBERT-BFD | ~420 Million | BFD (2.1 billion protein sequences) | 1024 |
| PepBERT | PepBERT | PepBERT-large-UniParc | ProtBERT-derived | Peptide-specific datasets | 1024 |

#### 6.3. Feature Fusion and Selection

To capture complementary sequence information associated with subcellular localization, individual descriptors and PLM embeddings were first evaluated independently and subsequently combined using a feature fusion strategy. The fused representations integrated various compositional and PLM embeddings of CPPs, providing a richer characterization of peptide sequences for ML-based localization prediction.

### 7. Classification Models

#### 7.1. Machine Learning (ML) Algorithms

Several ML methods were evaluated for CPP identification and localization prediction, including Decision Tree (DT), Random Forest (RF), Extra Trees (ET), Gradient Boosting, AdaBoost, Extreme Gradient Boosting (XGBoost), CatBoost, Logistic Regression (LR), and Quadratic Discriminant Analysis (QDA). Models were implemented using the Scikit-learn library and in-house Python scripts [61]. Hyperparameter optimization was performed using grid search within five-fold CV, with the area under the receiver operating characteristic curve (AUC) used as the optimization criterion.

#### 7.2. Deep Learning (DL) Algorithms

DL methods have been widely applied in bioinformatics owing to their ability to automatically learn complex representations from biological data [62]. Hence, to complement conventional ML approaches, four DL architectures were evaluated, including Artificial Neural Network (ANN), Deep Neural Network (DNN), One-Dimensional Convolutional Neural Network (1D-CNN), and Long Short-Term Memory (LSTM). All models were implemented using TensorFlow/Keras and trained using the Adam optimizer with binary cross-entropy loss, a batch size of 16, and 30 training epochs.

### 8. Ensemble approach for classification

To further enhance prediction performance, a hybrid framework integrating ML predictions with motif-based information derived from MERCI was developed. We collected five sets of motif-specific motifs for each location and utilized them to adjust the prediction probabilities. Motifs obtained with high accuracy were included in the hybrid prediction. A weighted scoring strategy was employed in which the presence of a localization-specific motif in the validation set contributed an additional score of “+0.5” to the class probabilities, whereas sequences without motif matches received a score of “0”. The final prediction score was calculated by combining the ML probability with the motif-derived score, thereby improving prediction reliability and interpretability. Similar hybrid strategies have already proven better performance in a range of bioinformatics prediction tools [2] [63] [64].

### 9. Performance Evaluation

Model performance was evaluated using Accuracy (Acc), Sensitivity (Sen), Specificity (Spec), Precision (Prec), F1-score, Matthews Correlation Coefficient (MCC), and Area Under the Receiver Operating Characteristic Curve (AUC). Performance metrics were computed for both the five-fold CV and the validation datasets. Definitions and mathematical formulations of these metrics have been described previously [65] [66]. The optimal decision threshold for each classifier was determined by selecting the point on the ROC curve where the difference between sensitivity and specificity was minimized, corresponding to the maximum Youden index.

### 10. Statistical Analysis

Statistical studies of AAC, DPC, and physicochemical properties were used to uncover distinguishing sequence characteristics associated with CPP activity and subcellular localization. Since the data did not meet the assumptions of normality and homoscedasticity, nonparametric statistical techniques were employed throughout the investigation. Differences between CPPs and non-CPPs were evaluated using the Mann-Whitney U test [67] using **scipy.stats.mannwhitneyu**, whereas comparisons among the five localization classes were performed using the Kruskal-Wallis test [68] using **scipy.stats.kruskal**. Cliff’s Delta ( and epsilon-squared statistics were used to calculate effect sizes for two-group and multi-group comparisons, respectively. To control for multiple hypothesis testing, p-values were adjusted using the Benjamini-Hochberg false discovery rate (FDR) correction **statsmodels.stats.multitest** module [69], and adjusted p-values < 0.05 were considered statistically significant. Dunn’s post hoc test [70] was performed using **scikit_posthocs.posthoc_dunn** was used to discover pairwise class differences in physicochemical parameters that differed significantly between localization classes.

### 11. Architecture of Web-server

A user-friendly web server, CPPLocPred (https://webs.iiitd.edu.in/raghava/cpplocpred/), was developed to facilitate the prediction of CPPs and their subcellular localization. The server was implemented using HTML5, CSS3, JavaScript, PHP, and responsive web templates, ensuring compatibility across desktop, tablet, and mobile devices. CPPLocPred provides modules for peptide prediction and motif searching, enabling users to submit peptide sequences and obtain CPP classification and localization predictions via an intuitive web interface.

## Results

### 1. Preliminary Sequence Analysis

#### 1.1. AAC Analysis

##### 1.1.1. Comparison between CPPs and Non-CPPs

To identify compositional differences related to cell-penetrating ability, AAC of CPPs and non-CPPs were compared using the Mann-Whitney U test followed by Cliff’s delta effect size analysis (Figure 3A and 3B). Several amino acids differed significantly, such as CPPs exhibiting higher proportions of positively charged residues, particularly R (24.14% v/s 5.87%) and K (14.75% v/s 6.71%), as well as increased frequencies of C (3.40% v/s 0.89%) and W (4.13% v/s 0.63%). In contrast, non-CPPs contained greater proportions of acidic residues, including D (4.99% v/s 1.46%) and E (6.83% v/s 2.25%), as well as A, I, S, T, V, and L. The significant residues have medium-to-large Cliff’s delta values, indicating substantial compositional differences between CPPs and non-CPPs (Supplementary Table S2.2).

**Figure 3:**
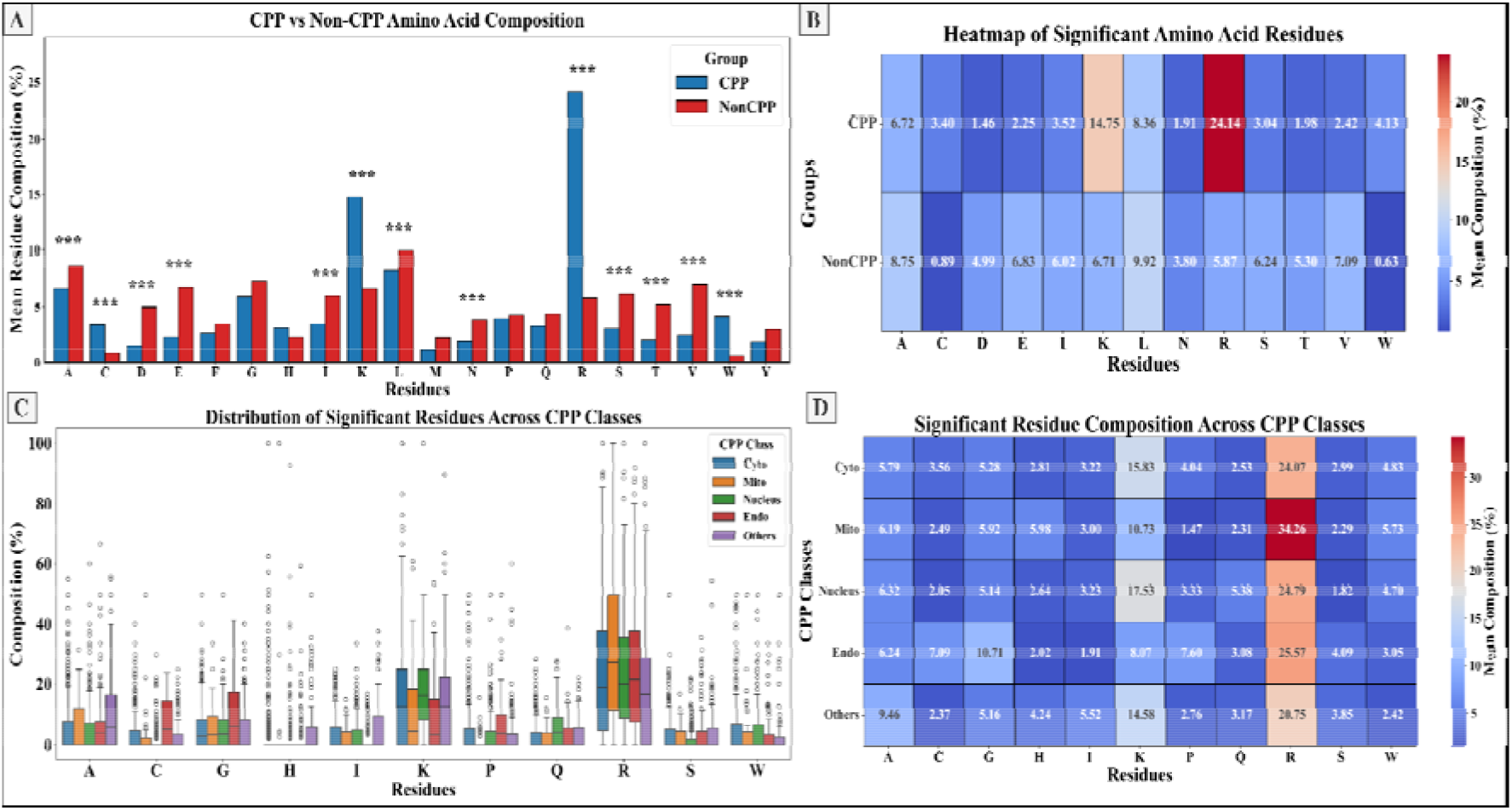
(A) Comparison of AAC between CPPs and non-CPPs. Asterisks indicate significant differences (*P* < 0.05). (B) Heatmap showing mean AAC across CPP localization classes. (C) Boxplots showing the distribution of significantly different amino acids across CPP localization classes. (D) Heatmap of residues significantly differentiating CPP classes.

##### 1.1.2. Localization-Specific AAC Analysis

The Kruskal-Wallis test was used to compare AACs among CPP localization classes (Supplementary Table S2.3). Figures 3C and 3D reveal distinct compositional trends among localization classes. Significant differences were found for several residues, including A, C, G, H, I, K, P, Q, R, S, and W, with the mean composition of all residues shown in Supplementary Table S2.1. Mitochondrial CPPs contained the highest proportions of R (34.26%) and combined basic residues (R, K, and H = 50.97%), but the lowest proportions of acidic residue content (D and E = 1.17%), P (1.46%), V (1.33%), and Q (1.47%). Nuclear CPPs were likewise richer in R (24.79%) residues and comparatively higher K (17.53%), E (3.08%), N (2.02%), M (1.52%), and Q (5.38%). In contrast, Endo_lysosomal CPPs showed increased frequencies of C (7.09%), G (10.71%), P (7.60%), S (4.09%), and V (3.30%), together with lower levels of K (8.07%) and L (5.93%) than Mitochondrial and Nuclear CPPs. Cytoplasmic CPPs have residue frequency patterns that are intermediate between Mitochondria, Nucleus, and Endo_lysosome classes. The Others localization class exhibited higher proportions of A (9.46%), L (10.35%), and I (5.52%), resulting in a relatively hydrophobic and aliphatic residue profile, indicating that they rely more on hydrophobic contacts than electrostatic interactions.

#### 1.2. DPC Analysis

##### 1.2.1. Comparison between CPPs and Non-CPPs

To examine sequence-order characteristics associated with cell penetration, DPC of CPPs and non-CPPs were compared. Figure 4B depicts the dipeptides that were significantly different between the two groups. CPPs were enriched in cationic dipeptides, including RR (11.90%), KK (4.64%), KR (2.15%), RK (2.16%), GR (1.54%), RQ (1.32%), RW (1.26%), and WR (1.09%). Among these, RR was the most abundant dipeptide and showed nearly 28-fold higher abundance in CPPs than in non-CPPs (11.90% vs. 0.43%). In contrast, non-CPPs had more dipeptides with acidic and hydrophobic residues, including VE, IV, DA, AE, VV, VA, EK, and SL. The mean value and P-value statistics for significant DPC revealed medium-to-large Cliff’s delta values, indicating significant DPC differences between CPPs and non-CPPs (Supplementary Tables S2.6 and S2.7).

**Figure 4:**
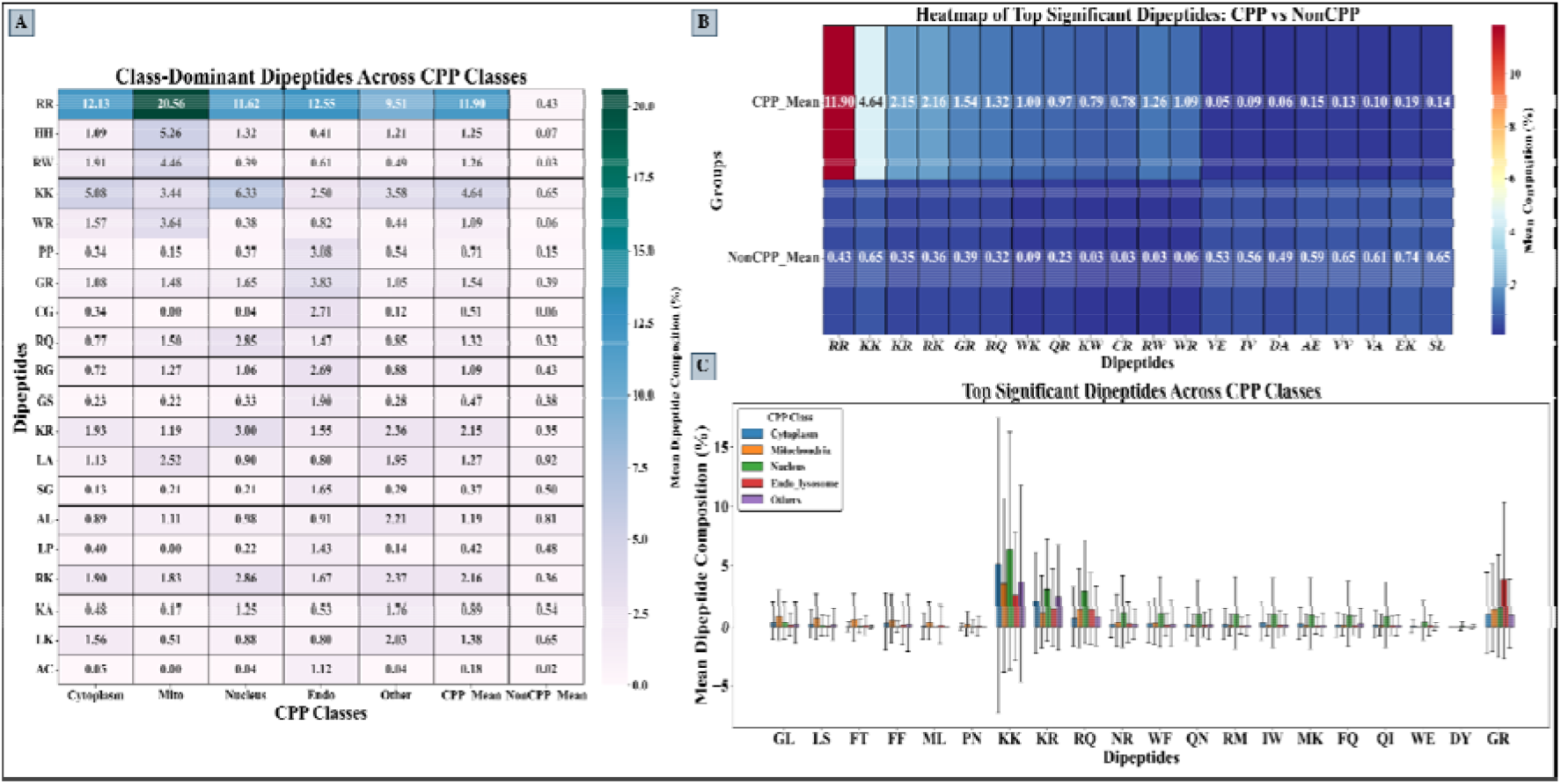
(A) Heatmap of dominant dipeptides across CPP localization classes. (B) Heatmap showing the top significant dipeptides that differentiate CPPs from non-CPPs. (C) Mean ± SD distribution of the top localization-associated dipeptides across CPP classes.

##### 1.2.2. Localization-Specific DPC Analysis

The DPCs were also compared among CPP localization classes using the Kruskal-Wallis test (Supplementary Table S2.8). Boxplot demonstrates that the DPC differed significantly amongst CPP localization classes (Figures 4A and 4C). Although RR was one of the most abundant dipeptides in all classes, its frequency varied significantly across subcellular locations, with Mitochondrial CPPs having the highest abundance (20.56%), followed by Endo_lysosomal (12.55%), Cytoplasmic (12.13%), Nuclear (11.62%), and Others CPPs (9.51%). Nuclear CPPs had the greatest frequencies of KK (6.33%), KR (3.00%), and RQ (2.85%), whereas Endo_lysosomal CPPs had higher abundances of GR (3.83%), PP (3.08%), CG (2.71%), RG (2.69%), and GS (1.90%). Aside from RR, Mitochondrial CPPs were distinguished by higher frequencies of HH (5.26%), RW (4.46%), and WR (3.64%). In contrast, the Others localization class exhibited higher proportions of LA (1.95%), AL (2.21%), KA (1.76%), LK (2.03%), in addition to RR, KK, KR and RK.

#### 1.3. Physicochemical Property Analysis

##### 1.3.1. Comparison between CPPs and Non-CPPs

To explore physicochemical characteristics associated with cell penetration, several sequence-derived properties were examined. Significant differences were observed across multiple properties, notably charge-related properties, and residue-group composition (Supplementary Table S2.4). CPPs exhibited substantially higher isoelectric points (pI) and net positive charges than non-CPPs, with mean pI values of 10.65 and 7.03 and mean net charges of 6.40 and 0.59, respectively (Figure 5A). Consistent with these observations, CPPs have a 3-fold greater fraction of basic residues (42.00%) than non-CPPs (14.82%). Similarly, CPPs have a very high instability index (103.67%). Non-CPPs had greater amounts of acidic residues (11.82% vs 3.71%), non-polar residues (49.15% vs 34.72%), aliphatic residues (39.15% vs 26.93%), hydrophobicity (-0.26% vs -1.38%), and neutral residues (73.36% vs 54.29%). CPPs also exhibited lower hydrophobicity values than non-CPPs (-1.38 vs. -0.26) (Figure 5C).

**Figure 5:**
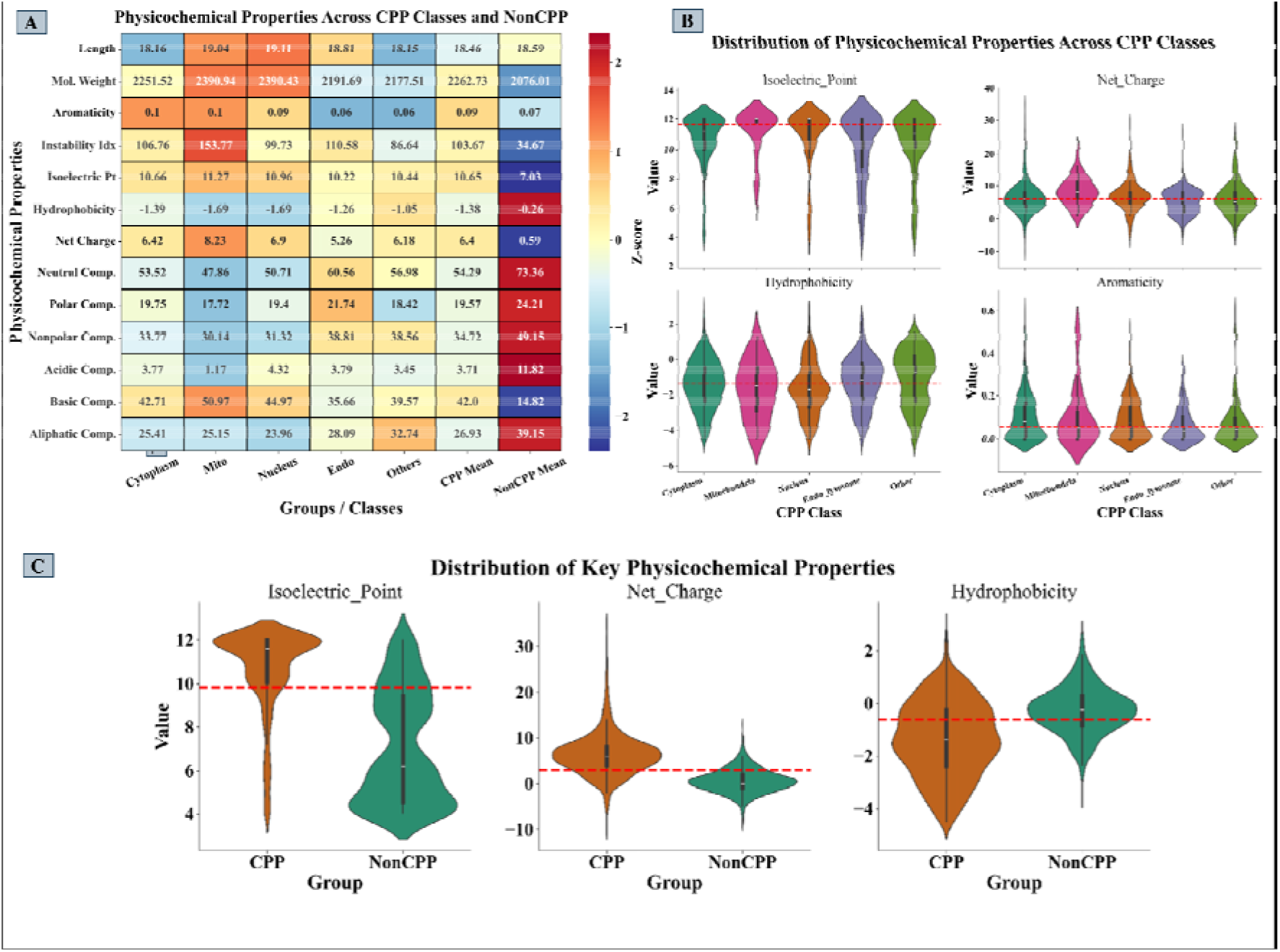
(A) Heatmap of mean physicochemical properties across CPP localization classes and non-CPPs. (B) Violin plots illustrate the distributions of physicochemical properties across CPP localization classes. (C) Violin plots showing the distributions of key physicochemical properties between CPPs and non-CPPs. The red-dashed line indicates the overall mean for each property across all peptides.

##### 1.3.2. Localization-Specific Physicochemical Properties

Comparison of physicochemical properties among localization classes revealed substantial diversity across subcellular targeting (Supplementary Table S2.5). Cytoplasmic CPPs displayed intermediate values, with a mean net charge of 6.42 and basic residue composition of 42.71%. Mitochondrial CPPs exhibited the highest isoelectric point (11.27), net charge (8.23), basic residue composition (50.97%) and instability index (153.77), suggesting increased structural flexibility. Nuclear CPPs also showed elevated pI (10.96), net charge (6.90), and basic residue content (44.97%). Endo_lysosomal CPPs were characterized by lower pI (10.22), net charge (5.26), and basic residue composition (35.66%) than the Mitochondrial and Nuclear classes. This class also exhibited the high polar residue content (21.74%) and non-polar residue composition (38.81%). The Others localization class showed the highest aliphatic residue composition (32.74%) and relatively low hydrophobicity (-1.05), indicating greater sequence diversity. Aromaticity varied relatively little between localization classes, from 0.06 to 0.10 (Figure 5B).

#### 1.4. Position-Specific Residue Analysis

##### 1.4.1. Comparison between CPPs and Non-CPPs

TSL Comparison of CPPs and non-CPPs revealed significant enrichment of positively charged residues, particularly R and K, across all positions (Figure 6A). Arginine was the most prominently enriched residue throughout the sequence. In contrast, acidic residues (D and E) and several hydrophobic residues were significantly depleted in CPPs relative to non-CPPs.

**Figure 6:**
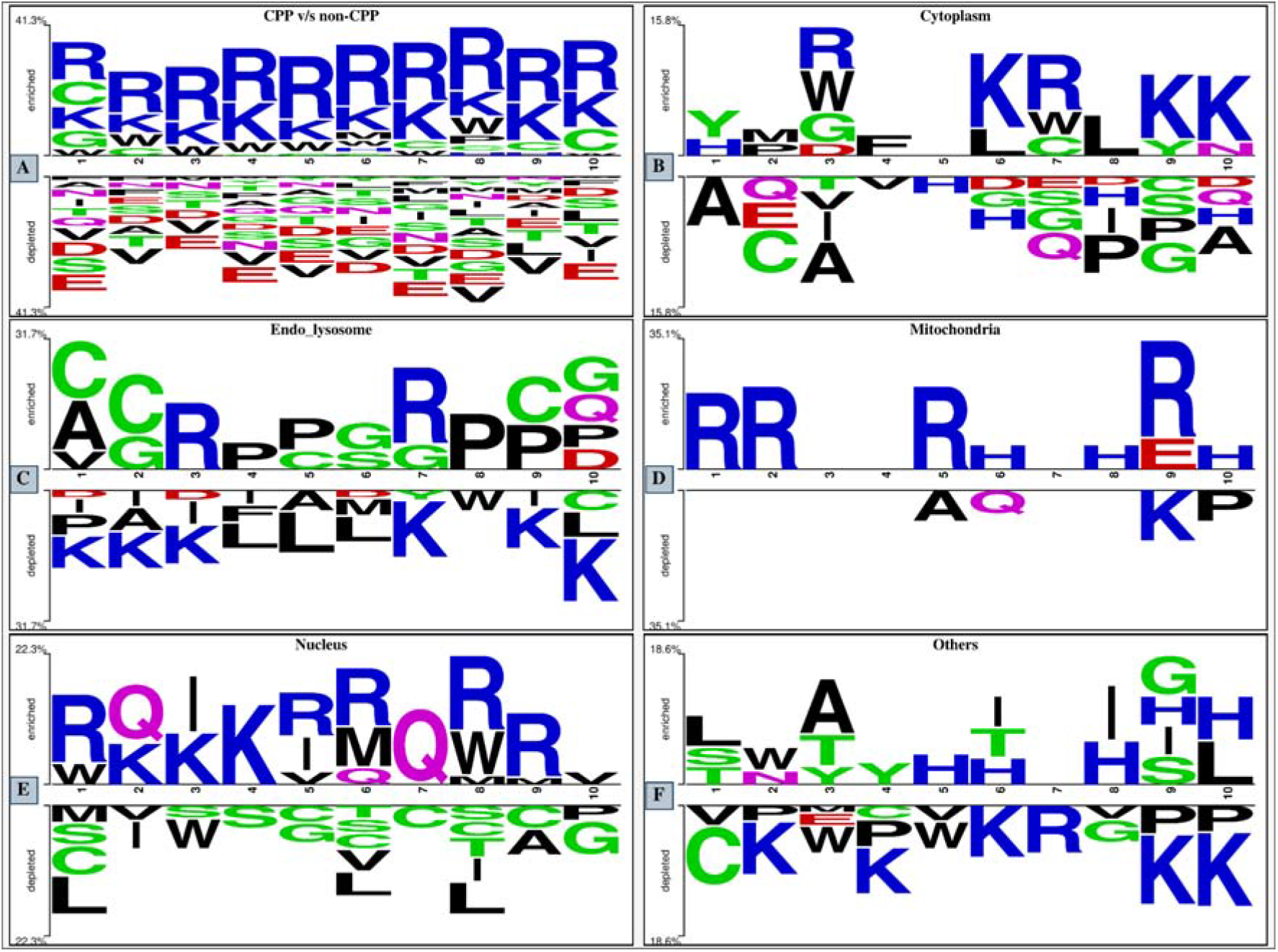
TSL analysis (A) CPPs and non-CPPs, (B) Cytoplasm, (C) Endo_lysosome, (D) Mitochondria, (E) Nucleus, (F) Others CPPs.

##### 1.4.2. Localization-Specific TSL Analysis

The localization-specific TSL analysis revealed distinct positioning residue patterns among CPP classes (Figures 6B, 6C, 6D, 6E & 6F). Cytoplasmic CPPs included more R and L residues, notably in the middle and C-terminal regions. R was enriched at the 3rd, 6th, and 7th positions, whereas K was enriched at the 6th, 9th, and 10th positions. Several acidic and polar residues were depleted. Mitochondrial CPPs showed a high enrichment of R residues at positions 1, 2, 5, and 9, as well as H at positions 6, 8, and 10. In contrast, K, A, Q, and P residues were decreased in many locations. Nuclear CPPs showed substantial enrichment of K and R residues across multiple positions, with successive enrichment from positions 3 to 9. Q was abundant at the 2nd, 6th, and 7th positions, while hydrophobic residues (L and V) and other residues (S, C, and G) were deficient. Endo_lysosome CPPs were represented by the enrichment of C, G, P, R, and A at several positions. In contrast, K and other basic residues were under-represented compared to the Mitochondrial and Nuclear classes. CPPs from the Others class had more hydrophobic and polar residues, such as A, T, H, S, G, L, and Y, in many positions. R and K residues were generally depleted.

### 2. Motif Analysis

Motif discovery using MERCI revealed several localization-associated sequence patterns across CPP classes. The “NONE” classification found specific amino acid motifs, whereas the BETTS-RUSSELL, KOOLMAN-ROHM, and RASMOL classifications identified motifs based on amino acid physicochemical groups. Supplementary Table S3 contains detailed motifs found using the mentioned classification schemes, showing distinct motif profiles for each CPP localization class. Cytoplasmic CPPs were characterized with motifs including KKLLK, LKKLL, RWKC, WKC, and WRWKC. Mitochondrial CPPs included motifs such as FT, FF, KIK, MLS, RLL, and polyarginine-rich sequences, such as RRRRRRRR. Nuclear CPPs exhibited motifs such as KKGG, AKKA, LY, DY, ELW, and GQ. Endo_lysosomal CPPs revealed enrichment of motifs containing cysteine and glycine residues, including CGR, GCG, RGR, ACR, SCG, and CGRK, whereas the Others class was related to motifs such as AGYLLG, GYLLG, LII, YLLG, and AGYLLGK. The physicochemical classification systems indicated different motif distributions across localization classes. Mitochondrial and Nuclear CPPs had a greater frequency of positively charged and hydrophobic residue groups, whereas Endo_lysosomal CPPs had motifs richer in G-, C-, and other polar residue groups. In contrast, the Others class had a greater abundance of hydrophobic and aliphatic residue-associated motifs.

### 3. Performance of ML Classifiers for CPP prediction

In this study, 16 sequence-derived compositional features were evaluated using multiple ML classifiers to distinguish CPPs from non-CPPs. The performance of the ET classifier on the top feature on the validation dataset is summarized in Table 4, while complete results are provided in Supplementary Table S4. Among the evaluated features, AAC achieved the best overall performance, yielding an AUC of 0.979, an MCC of 0.847, an accuracy of 92.3%, a sensitivity of 93.4%, and a specificity of 91.2%. Comparable performance was also observed for PAAC (AUC = 0.978), QSO (AUC = 0.976), APAAC (AUC = 0.976), and SER (AUC = 0.974). Based on its superior predictive performance on the validation dataset, the AAC-ET model was selected as the final CPP predictor, and the resulting threshold of 0.44 was subsequently used for independent testing and deployment.

**Table 4:**
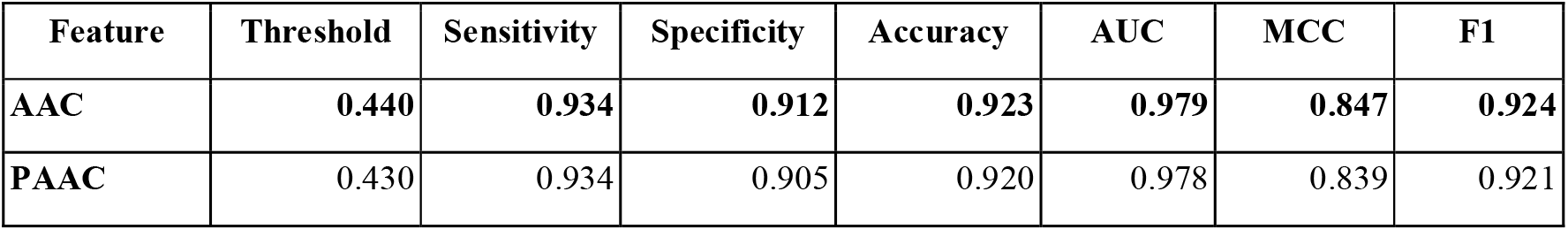
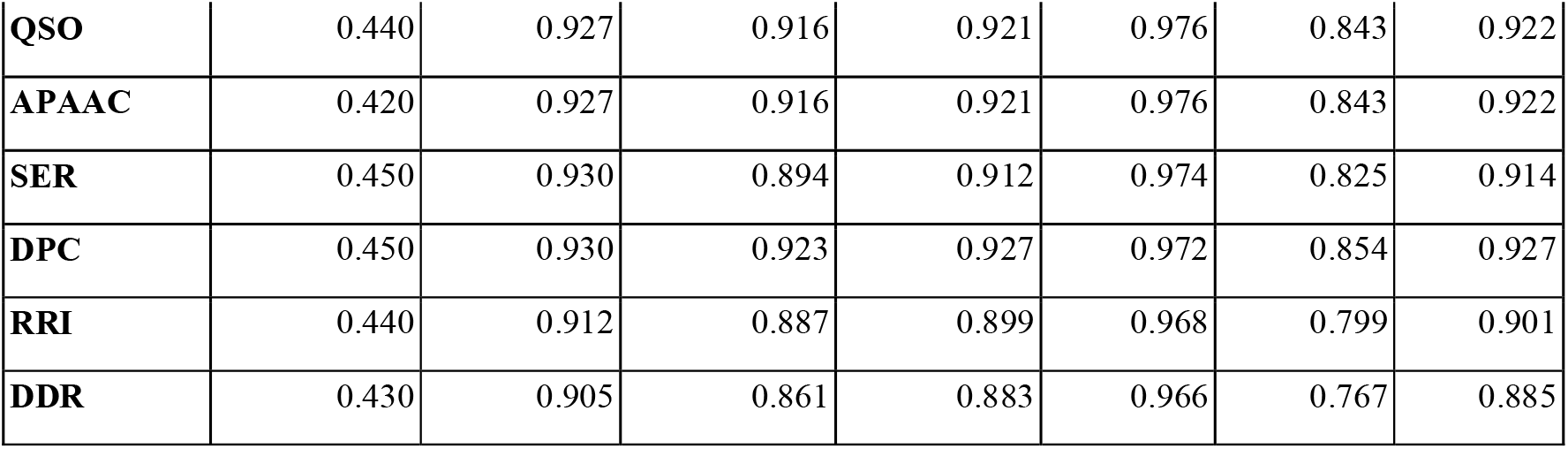
Performance of the ET model on the top compositional features on the validation dataset for CPP prediction.

### 4. Benchmarking CPP prediction model on existing method

As shown in Table 5, we benchmarked the best-performing CPP classifier against existing methods, using a separate independent dataset comprising 1,516 peptides (758 CPPs and 758 non-CPPs) to assess the robustness and generalization ability. Several additional CPP prediction methods were evaluated for comparison. However, many of these methods either lack publicly available standalone implementations or pretrained models, are no longer maintained, or require retraining from source without providing inference-ready models. Consequently, only methods that could be executed reproducibly (MLCPP [28], PerseuCPP [37]and pLM4CPPs [71]) were taken into the comparative analysis. On the complete independent dataset, the CPP classifier of CPPLocPred obtained the best performance among all approaches, with an accuracy of 89.12%, and an AUC of 0.953, outperforming MLCPP (accuracy: 83.97%, AUC: 0.9213), PerseuCPP (accuracy: 86.07%, AUC: 0.9289) and pLM4CPPs (accuracy: 84.83%). Similar trends were observed across all CD-HIT filtered datasets, where CPPLocPred consistently maintained superior or comparable performance.

**Table 5:** Performance comparison of the existing tool and our CPP prediction on the separate independent set.

| CD-Hit<br>(Seq count) | Our Method |  | MLCPP |  | PerseuCPP |  | pLM4CPPs (ESM-640) |  |
| --- | --- | --- | --- | --- | --- | --- | --- | --- |
|  | AUC | Accuracy | AUC | Accuracy | AUC | Accuracy | AUC | Accuracy |
| Full set (1516 seq) | **0.953** | **0.8912** | 0.9213 | 0.8397 | 0.9289 | 0.8607 | N/A | 0.8483 |
| 90% (890 seq) | **0.9422** | 0.8753 | 0.9191 | 0.8517 | 0.9302 | 0.8719 | N/A | 0.8517 |
| 80% (770 seq) | **0.9284** | 0.8571 | 0.9069 | 0.8416 | 0.9202 | 0.8636 | N/A | 0.8442 |
| 70% (680 seq) | **0.9189** | 0.8471 | 0.898 | 0.8353 | 0.9179 | 0.8588 | N/A | 0.8338 |
| 60% (541 seq) | 0.8978 | 0.8262 | 0.8803 | 0.8226 | **0.8988** | 0.8355 | N/A | 0.8152 |
| 50% (294 seq) | **0.8908** | 0.8333 | 0.8872 | 0.8435 | 0.8846 | 0.8299 | N/A | 0.8027 |
| 40% (115 seq) | 0.8582 | 0.8957 | **0.9** | 0.8783 | 0.852 | 0.8696 | N/A | 0.8435 |

### 5. Subcellular Localization Prediction Models

#### 5.1. Performance of ML- and DL-based Models Using Compositional Features

To establish baseline localization prediction models, 16 compositional features were evaluated using multiple ML and DL algorithms and the best-performing features on respective models were depicted in Tables 6 and 7. Five independent binary classifiers corresponding to Cytoplasm, Mitochondria, Nucleus, Endo_lysosome, and Others localization classes were developed using an OvR strategy. The compositional features that showed the best overall performance after ML classifiers, such as AAC, DPC, CETD, DDR, RRI, QSO, and the fused ALLCOMP, were subsequently selected for DL model development and comparative evaluation. Complete performance for all models is provided in the Supplementary Tables S5, S6, S7, S8, S9 and S10.

**Table 6:** Validation AUC of the best-performing ML model and compositional features for subcellular localization prediction.

| Feature Set | Cytoplasm | Mitochondria | Nucleus | Endo_lyosome | Others | Mean AUC |
| --- | --- | --- | --- | --- | --- | --- |
| AAC | 0.800 (RF) | 0.845 (ET) | 0.805 (ET) | 0.797 (CB) | 0.782 (RF) | 0.806 |
| DDR | 0.814 (CB) | 0.970 (CB) | 0.775 (CB) | 0.791 (RF) | 0.798 (CB) | **0.830** |
| DPC | 0.806 (RF) | 0.860 (RF) | 0.804 (RF) | 0.811 (XGB) | 0.809 (ET) | 0.818 |
| CETD | 0.811 (XGB) | 0.731 (AB) | 0.797 (XGB) | 0.791 (RF) | 0.814 (XGB) | 0.789 |
| QSO | 0.818 (RF) | 0.886 (CB) | 0.794 (ET) | 0.797 (ET) | 0.790 (ET) | 0.817 |
| RRI | 0.828 (RF) | 0.917 (CB) | 0.784 (ET) | 0.800 (RF) | 0.793 (RF) | 0.824 |
| Combined (1165) | 0.841 (RF) | 0.833 (LR) | 0.815 (ET) | 0.815 (ET) | 0.837 (RF) | 0.828 |

#RF, Random Forest; ET, Extra Trees; CB, CatBoost; XGB, XGBoost; LR, Logistic Regression; AB, AdaBoost.

**Table 7:** Validation AUC of the best-performing DL model and compositional features for subcellular localization prediction.

| Feature Set | Cytoplasm | Mitochondria | Nucleus | Endo_lysome | Others | Mean AUC |
| --- | --- | --- | --- | --- | --- | --- |
| AAC | 0.777 (DNN) | 0.833 (DNN) | 0.749 (DNN) | 0.787 (ANN) | 0.743 (DNN) | 0.778 |
| DDR | 0.786 (DNN) | 0.833 (LSTM) | 0.764 (DNN) | 0.799 (DNN) | 0.706 (DNN) | 0.778 |
| DPC | 0.797 (DNN) | 0.833 (ANN) | 0.744 (DNN) | 0.786 (DNN) | 0.783 (DNN) | 0.789 |
| CETD | 0.754 (ANN) | 0.758 (LSTM) | 0.701 (ANN) | 0.791 (DNN) | 0.740 (DNN) | 0.749 |
| QSO | 0.759 (DNN) | 0.856 (DNN) | 0.745 (ANN) | 0.812 (DNN) | 0.707 (DNN) | 0.776 |
| RRI | 0.807 (ANN) | 0.924 (DNN) | 0.716 (DNN) | 0.762 (DNN) | 0.725 (ANN) | 0.787 |
| Combined (1165) | 0.807 (ANN) | **0.939 (ANN)** | 0.718 (ANN) | 0.769 (ANN) | 0.762 (ANN) | **0.799** |

#ANN, Artificial Neural Network; DNN, Deep Neural Network; LSTM, Long Short-Term Memory.

#### 5.2. Performance of PLM Embeddings

ML models trained on PLM embeddings have been widely used in studies to investigate the contribution of contextual sequence information [72] [73]. In this study, we have utilized embeddings from pretrained ProtBERT, PepBERT, ESM2-8M, ESM2-35M, and ESM2-150M models. Fine-tuning was not performed because the localization datasets were relatively small (114-1326 samples per binary classification task), which could increase the risk of overfitting when training large transformer models. Instead, the extracted pretrained embeddings were used as input features for ML classifiers for model development. As shown in Table 8, performance varied among embedding models, with ESM2-derived embeddings generally producing the highest predictive performance across localization classes. PepBERT embeddings also demonstrated competitive performance, whereas ProtBERT embeddings performed comparatively worse. Detailed performance statistics for all embedding-based models for each localization class are provided in the Supplementary Table S11.

**Table 8:** Validation AUC of best-performing ML models on PLM embeddings for subcellular localization prediction.

| Embedding | Cytoplasm | Mitochondria | Nucleus | Endo_lysome | Others | Mean AUC |
| --- | --- | --- | --- | --- | --- | --- |
| ESM2_8M | 0.819 (CB) | 0.909 (GB) | 0.792 (CB) | 0.823 (XGB) | 0.737 (CB) | **0.816** |
| ESM2_35M | 0.812 (RF) | 0.848 (GB) | 0.806 (CB) | 0.817 (ET) | 0.748 (XGB) | 0.806 |
| ESM2_150M | 0.803 (ET) | 0.848 (XGB) | 0.822 (CB) | 0.793 (RF) | 0.791 (XGB) | 0.811 |
| PepBERT | 0.827 (ET) | 0.860 (ET) | 0.744 (CB) | 0.809 (LR) | 0.768 (CB) | 0.802 |
| ProtBERT | 0.783 (ET) | 0.826 (DT) | 0.785 (ET) | 0.739 (RF) | 0.748 (ET) | 0.776 |
| Combined | 0.811 (ET) | 0.886 (GB) | 0.818 (XGB) | 0.814 (LR) | 0.749 (RF) | 0.816 |

#RF, Random Forest; ET, Extra Trees; CB, CatBoost; XGB, XGBoost; GB, GradientBoost; LR, Logistic Regression.

#### 5.3. Selection of the Final Localization Prediction Model

To further improve subcellular localization prediction, feature fusion models were developed by combining the best-performing compositional and PLM embeddings. Multiple fusion strategies were evaluated, and their corresponding performance statistics are provided in the Supplementary Table S12. Lastly, to identify the optimal localization predictor, all feature-classifier combinations were compared using the mean validation AUC and MCC across the five localization classifiers as reported in Supplementary Table S13. This model-agnostic evaluation provided an overall assessment of predictive performance while minimizing bias associated with individual localization classes. Among all evaluated combinations, the 6F_526 with the ET model achieved the highest validation AUC of 0.831 ± 0.034, as shown in Table 9. However, this model required a relatively large feature vector consisting of 526 features. In contrast, the DDR with the CatBoost model achieved a nearly identical validation AUC of 0.828 ± 0.081 with a higher validation MCC (0.506 ± 0.232 vs 0.473 ± 0.031), indicating superior overall classification quality across localization classes. Given the negligible difference in AUC, the superior MCC, and the substantially lower feature dimensionality (20 features), the DDR-CatBoost model was selected as the final subcellular localization predictor.

**Table 9:** Performance comparison of the Best Feature-Classifier combinations based on mean AUC and MCC across the five localization classifiers.

| Feature | Model | Train_AUC | Test_AUC | Train_MCC | Test_MCC |
| --- | --- | --- | --- | --- | --- |
| 6F_526 | ET | $0.764 \pm 0.027$ | $0.831 \pm 0.034$ | $0.438 \pm 0.069$ | $0.473 \pm 0.031$ |
| **DDR** | **CatBoost** | **$0.713 \pm 0.041$** | **$0.828 \pm 0.081$** | **$0.376 \pm 0.059$** | **$0.506 \pm 0.232$** |
| 5F_126 | ET | $0.754 \pm 0.025$ | $0.827 \pm 0.036$ | $0.429 \pm 0.057$ | $0.494 \pm 0.117$ |
| DDR_RRI | CatBoost | $0.728 \pm 0.041$ | $0.826 \pm 0.061$ | $0.364 \pm 0.047$ | $0.537 \pm 0.134$ |
| DDR_ESM8M_340 | ET | $0.750 \pm 0.018$ | $0.822 \pm 0.061$ | $0.419 \pm 0.046$ | $0.443 \pm 0.114$ |
| ALL COMP _1165 | ET | $0.771 \pm 0.021$ | $0.821 \pm 0.013$ | $0.451 \pm 0.064$ | $0.505 \pm 0.040$ |
| RRI | RF | $0.745 \pm 0.033$ | $0.816 \pm 0.051$ | $0.388 \pm 0.051$ | $0.463 \pm 0.124$ |
| QSO | ET | $0.750 \pm 0.030$ | $0.810 \pm 0.027$ | $0.423 \pm 0.054$ | $0.403 \pm 0.059$ |
| APAAC | ET | $0.755 \pm 0.022$ | $0.809 \pm 0.023$ | $0.421 \pm 0.043$ | $0.446 \pm 0.068$ |
| DPC | RF | $0.746 \pm 0.032$ | $0.807 \pm 0.032$ | $0.420 \pm 0.072$ | $0.455 \pm 0.075$ |

Additionally, to enhance robust prediction performance further for the subcellular localization classifiers, we implemented an ensemble strategy that integrates the best-performing subcellular localization predictor with an alignment-based approach using Motif search. Motifs exhibiting high coverage and predictive accuracy across different parameter combinations were incorporated into the prediction process using a weighted scoring strategy. However, the performance is roughly the same as that of the ML model and doesn’t improve significantly. Detailed performance of the hybrid models is provided in Supplementary Table S15.

### 6. Web Server Implementation

To facilitate the scientific community, the final prediction models were implemented in a user-friendly web server, CPPLocPred at https://webs.iiitd.edu.in/raghava/cpplocpred/. The web server integrates three major modules: prediction, design, and motif scan. The ‘Prediction module’ enables users to screen and classify input sequences as CPPs or non-CPPs, and then determine subcellular location using a threshold, with probabilities displayed. In this module, Users can submit single or multiple peptide sequences in FASTA format and obtain prediction scores, probabilities, and subcellular location classes. In the “Design module”, the user can design new variants of different subcellular localization CPPs. In addition, a “Motif Scan” module was developed to identify localization-associated motifs discovered using MERCI. It scans the user-provided query sequence for patterns. In addition, we provide a CPPLocPred standalone Python package, accessible via the web server’s “Download” section, to enable users to identify CPP and their intracellular locations at a larger scale.

## Discussion

CPPLocPred is a considerable improvement over current CPP prediction methods since it integrates subcellular localization prediction into a unified framework. Most currently available predictors rely solely on identifying CPPs from non-CPPs, offering little information about the intracellular fate of transported peptides. The hierarchical architecture developed in this study addresses this challenge by first identifying CPPs and then predicting their subcellular localization, resembling the normal biological progression of peptide absorption and transport.

The preliminary sequence analyses conducted in this study demonstrated that intracellular localization is associated with different sequence signatures that extend beyond general cell-penetrating ability. The most prominent feature was the strong enrichment of R and K residues in CPPs. The presence of positive charges on these residues provides opportunities for electrostatic interactions with negatively charged glycosaminoglycans, proteoglycans, and phospholipid membranes, which appear to be among the early steps in CPP uptake [17] [74]. Among those residues, R is the most crucial, since the guanidinium side chain allows R to form multiple hydrogen bonds and electrostatic interactions simultaneously, providing greater effectiveness in membrane translocation than K-rich peptides [16] [15]. Consistent with previous studies, a significant depletion in acidic residues in CPPs indicated that excessive negative charges may interfere with membrane binding and cell uptake [14].

Although AAC, DPC, physicochemical properties, positional residue preferences, and motif analyses evaluate peptide sequences from different perspectives, all approaches converge on a common observation that different localization classes possess characteristic sequence architectures. A unique cationic amphipathic profile for Mitochondrial CPPs was observed with the presence of higher frequencies of K, R, H, W, L; the highest frequency of RR dipeptides; higher net charge, isoelectric point, and content of basic residues and reduced content of acidic residues among all classes. The coexistence of both positively charged and hydrophobic/aromatic residues indicates an amphipathic structure typical of MTPs [18], which can assist electrostatic attraction to negatively charged mitochondrial membranes while facilitating association and translocation through hydrophobic contacts, in accordance with the properties of canonical mitochondrial targeting sequences [52] [75]. Similarly, Nuclear CPPs displayed elevated K and R content, increased frequency of KK- and KR-containing dipeptides, a positional basic residues cluster, and motifs resembling well-established NLS, recognized by importin-mediated transport pathways, in the sequences of Nuclear CPPs [51] [19].

Cytoplasmic CPPs displayed comparatively balanced residue compositions without extreme enrichment of specific amino acids, suggesting a more general CPP feature optimized for cytoplasmic distribution rather than organelle-specific targeting. On the other hand, the Endo_lysosomal CPPs exhibited a distinct profile from Mitochondrial and Nuclear CPPs, characterized by lower cationicity and an enrichment of C, G, P with a greater abundance of CG, GR and PP-rich di- and higher peptides. These features can enable greater structural flexibility, thus enhancing membrane remodelling and vesicle membrane interaction [76]. Moreover, the higher content of nonpolar residues can increase the tendency for membrane interaction and disruption. Previous studies have shown that a higher content of flexible amphipathic peptides containing G and P facilitates intracellular translocation and endosomal escape by transiently perturbing the membrane rather than relying solely on electrostatic binding [77] [78]. The presence of C-containing motifs might further contribute to intracellular targeting via thiol-based interactions [79].

Peptides assigned to the Others class had a greater number of hydrophobic and aliphatic residues and hinted at less specific association with the cellular membrane and less restricted interaction with lipids, as compared to targeted organelle-specific electrostatic interactions. This aliphatic character would facilitate better partitioning into cell membranes and a looser subcellular distribution behaviour. Collectively, these observations indicate that subcellular targeting is determined not only by overall positive charge but also by residue arrangement, positional preferences, and motif architecture. As a result, peptides with comparable residue compositions may display diverse localization behaviours due to changes in sequence organization and local residue clustering. The observed consistency across independent analytical methodologies raises confidence that the detected sequence signatures represent true biological factors of CPP localization rather than artefacts of a specific analytical technology [80] [81].

The unambiguous, localization-specific signature features discovered in preliminary analyses provide a strong biological foundation for the development of ML models. Given the superior performance observed for the classification of CPPs (AUC=0.979 and MCC=0.847 for AAC-Extra Trees model), it is probable that CPPs possess highly distinguishable compositional characteristics relative to non-CPPs. On the other hand, localization prediction was expected to be considerably more difficult since all localization classes consist of known biologically active CPPs sharing similar cell-penetrating features. As a result, accurate localization prediction relies on recognising subtle sequence changes linked with intracellular targeting rather than just finding generic CPP features. The ML models’ high accuracy in differentiation of localization classes thus indirectly supports the use of localization-associated signatures as biologically relevant.

In line with recent studies, ESM2 and PepBERT embeddings displayed strong predictive performance. However, in this study, PLM embeddings did not markedly outperform the best compositional descriptors. Indeed, AAC, DPC, DDR, RRI, and QSO exhibited predictive performance similar to, and in many cases better than, embedding-based features. This indicates that the major features regulating CPP localization are, to a significant extent, captured by AAC and sequence order information, and hence can be captured by more conventional descriptors. Still, the fact that ESM2 and PepBERT provide very competitive results implies that contextual sequence information has additional predictive value and might become more important with the increase in size of future localization datasets.

The ensemble models, ET and CatBoost classifiers, achieved consistent, significantly higher performance than other ML and DL algorithms across different representations, likely owing to their ability to efficiently handle nonlinear dependencies whilst simultaneously avoiding overfitting to noisy variables. Despite having the potential for greater discriminative ability, DL models failed to outperform classical ML models considerably, likely because the localization datasets are small, particularly those of minority groups of peptides such as Mitochondrial CPPs. Similar trends have been observed in other peptide prediction experiments, where carefully engineered sequence descriptors combined with ensemble learning algorithms yielded surprisingly competitive results compared to DL-based methods. Although the 6F526-ET model resulted in the best validation AUC (0.831), we selected the DDR-CatBoost as the final prediction model, as it provides a comparable AUC (0.828) as well as higher MCC (0.506) with a significant feature dimension reduction from 526 features of the 6F526-ET representation to only 20 features for DDR-CatBoost. Small degradation in predictive performance, but the significant reduction in the number of features offers several benefits, such as lower computational complexity, reduced memory requirements, better interpretability, and decreased risk of overfitting. The higher MCC also implies that the predictions given by DDR-CatBoost were more balanced across the various localization classes, including minority categories. Further, fine-tuning of CatBoost improved both the MCC and the accuracy of CatBoost for Cytoplasm, Nucleus, Endo_lysosome and Others localization classes, but for the Mitochondrial dataset, the best performances were obtained with the default parameters of CatBoost, likely because of a smaller number of samples and a higher propensity for overfitting as depicted in Supplementary Table S14. To further improve prediction performance, motif information derived using MERCI was incorporated into ML prediction models via a hybrid approach. However, the hybrid motif-assisted strategy did not significantly improve prediction accuracy beyond the DDR-CatBoost model alone; the identified motifs provided valuable biological insights and independently validated the sequence signatures identified in other analyses.

The final AAC-ET and DDR-CatBoost models were further combined in a two-step prediction pipeline, which firstly distinguishes CPPs from non-CPPs and secondly assigns predicted CPPs into one of the five subcellular localization categories and were implemented by a user-friendly CPPLocPred web server and standalone package. This will also provide insights into the rational design of CPP-mediated delivery systems with targeted delivery to specific subcellular locations, as well as further studies on peptide-mediated cellular uptake and transport. Overall, CPPLocPred provided a practical resource for studying CPP biology and designing peptide-based therapeutic delivery systems.

## Limitations and Future Perspectives

Despite promising analytical results and ML performance obtained in this work, some limitations need to be addressed: 1) Datasets for certain subcellular locations are relatively undersampled (like Mitochondrial CPPs), which may compromise the capability of the DL architecture to fully exploit complex sequence patterns. 2) Only 5 major subcellular localization were considered in the present study, while a lot of CPPs undergo trafficking among multiple cellular compartments dynamically. 3) PLMs help performance, but an end-to-end fine-tuning will be attainable with a larger experimentally validated localization data set size. Future developments may include more subcellular locations, prediction of multilabel localization, and experimentally derived structures. Incorporating peptide structural features and a sophisticated transformer architecture might help enhance prediction accuracy and reveal molecular features of CPP cellular trafficking.

## Conclusion

CPPLocPred, a ML framework for the prediction of CPPs and their subcellular localization. The system combines sequence-derived features, advanced ML algorithms, and motif-based interpretation to provide accurate and biologically meaningful predictions. The integrated system provided an efficient and effective computational tool for identifying new candidate CPPs with intracellular specificity, thereby promoting further studies on peptide drugs and targeted drug delivery. CPPLocPred provides a valuable resource for peptide therapeutics, intracellular delivery, and the rational design of compartment-targeting CPPs.

## Data Availability

The source code and datasets generated for this study can be accessed on the “CPPLocPred” web server at https://webs.iiitd.edu.in/raghava/cpplocpred/download.php and are publicly available on GitHub at https://github.com/namanm04/CPPLocPred.

## Amino Acid Abbreviations

A: Alanine
R: Arginine
N: Asparagine
D: Aspartic Acid
C: Cysteine
E: Glutamic Acid
Q: Glutamine
G: Glycine
H: Histidine
I: Isoleucine
L: Leucine
K: Lysine
M: Methionine
F: Phenylalanine
P: Proline
S: Serine
T: Threonine
W: Tryptophan
Y: Tyrosine
V: Valine

## Authors’ contributions

The dataset collection, processing, algorithm implementation, and prediction model development were done by NB. NB and GPSR analysed the results. NKM developed the web server. NB drafted the manuscript, and NB, NKM, and GPSR contributed to its writing and revision. GPSR conceived and coordinated the project. The final manuscript was read and approved by all authors.

## Acknowledgements

We would like to thank the Council of Scientific & Industrial Research (CSIR), Department of Biotechnology (DBT, India), for fellowships and financial support, and the Department of Computational Biology, IIITD, New Delhi, for infrastructure and facilities. We also want to acknowledge that figures were created using <u>Draw.io</u>.

